# Nrg1 regulates cortical wiring and motor recovery upon traumatic injury

**DOI:** 10.1101/2025.11.18.689009

**Authors:** Ana González-Manteiga, Carmen Navarro-González, Ángela Rodríguez-Prieto, Maria Vittoria Zavaglia, Pietro Fazzari

## Abstract

Traumatic brain injury (TBI) is a leading cause of disability, yet molecular mechanisms supporting cortical repair remain poorly defined. While Neuregulin-1 (Nrg1) is essential for cortical development, its role in traumatic cortical injury in adults is unclear. To circumvent developmental confounds, we used an inducible conditional Nrg1 knockout in the adult mouse and subjected it to controlled cortical damage (CCD) in the motor cortex. We combined high-resolution adeno-associated viral tracing of callosal projections with comprehensive behavioral, histological, and molecular analyses. Nrg1 deletion led to significant impairments in structural connectivity and long-term motor recovery, which were markedly exacerbated in aged mice, indicating a critical role for Nrg1 in adult cortical repair. Mechanistically, our data indicated that Nrg1 promoted this plasticity through its intracellular domain (ICD) signaling, acting cell-autonomously to enhance axonal outgrowth. Furthermore, loss of Nrg1 was associated with altered structure of perineuronal-nets (PNN) and increased neuroinflammation at the lesion site. Our findings identify endogenous Nrg1 as a key regulator of structural preservation and functional recovery, highlighting the Nrg1 signaling as a potential target to enhance cortical plasticity after trauma.

## Introduction

Traumatic brain injury (TBI) is a major cause of adult disability, imposing a substantial socioeconomic burden and limiting patients’ quality of life. Despite intense research, effective molecular strategies to promote neural repair remain scarce. The pathophysiology of TBI unfolds in distinct phases: an early wave of excitotoxicity and neuronal death, followed by inflammation and axonal retraction, and later a period of limited structural plasticity (Galgano et al. 2017; Joy and Carmichael 2021; Moskowitz et al. 2010). Although inflammation contributes to repair, excessive or prolonged activation worsens damage (Agirman et al. 2017; Anrather and Iadecola 2016). In the subacute and chronic phases, neuronal circuits attempt to rewire, but regeneration remains restricted by incomplete reactivation of developmental programs and by inhibitory extracellular cues (Curcio and Bradke 2018).

Neuregulin-1 (Nrg1) is a trophic factor that regulates key developmental processes, including neuronal migration, neurite outgrowth, myelination, and excitatory/inhibitory balance (Mei and Nave 2014; Rodríguez-Prieto et al. 2024). Altered Nrg1 forward signaling via ErbB4 signaling has been implicated in psychiatric disorders, but accumulating evidence also suggests a neuroprotective role in CNS injury (Kataria et al. 2019; Wu et al. 2015). Moreover, Nrg1 undergoes regulated intramembrane proteolysis by γ-secretase, releasing the intracellular domain (Nrg1-ICD), which translocates to the nucleus and modulates gene expression (Bao et al. 2003; Chen et al. 2010; Fazzari et al. 2014; Rodríguez-Prieto et al. 2024). This pathway has been linked to neuronal survival under stress (Navarro-González et al. 2019) and axonal growth during development (Rodríguez-Prieto et al. 2024), suggesting it could also contribute to repair after injury.

Most studies to date have focused on administration of exogenous Nrg1 after stroke, which improves recovery and reduces tissue damage (Alizadeh et al. 2017; Guan et al. 2015; Deng et al. 2019). However, the physiological role of endogenous Nrg1 in traumatic injury, particularly its non-canonical intracellular signaling, remains poorly defined. So far, the relevance of endogenous Nrg1/ErbB4 signaling has only been investigated in constitutive Nrg1 heterozygous mice (Noll et al., 2019) or in mice with conditional deletion of ErbB4 in parvalbumin (PV)-positive interneurons (Guan et al., 2015; Deng et al., 2019). Both models display marked developmental alterations (Fazzari et al., 2010; del Pino et al., 2013; Navarro-González et al., 2021, representing an important caveat for interpreting results in the context of adult injury.

To overcome these limitations, we generated an inducible Nrg1-deficient mouse line, enabling deletion of Nrg1 in the adult brain and thus avoiding developmental confounds. Taking advantage of this new tool, we investigated the physiological role of endogenous Nrg1 in the context of traumatic cortical injury. To do so, we combined complementary *in vitro* and *in vivo* approaches. First, we examined whether activation of intracellular Nrg1 signaling is sufficient to promote axonal regeneration in cultured cortical neurons. Second, to test the requirement for Nrg1 *in vivo*, we used a conditional knockout strategy in a refined model of controlled cortical damage (CCD) that allows reproducible injury to the motor cortex while preserving surrounding tissue (González-Manteiga et al. 2022). This model enabled us to analyze circuit remodeling with high anatomical resolution. We focused on interhemispheric callosal projections, which are crucial for motor coordination and represent a major site of cortical rewiring after injury (Brown, et al. 2009; Takatsuru, et al. 2009; Rehme, et al. 2011; Empl, et al. 2022; Chovsepian, et al. 2022). In parallel, we assessed the impact of Nrg1 loss on neuroinflammation and on the integrity of perineuronal nets surrounding parvalbumin-positive interneurons, a key inhibitory population tightly regulated by Nrg1/ErbB4 signaling. Finally, we evaluated motor performance in young and mature mice to determine whether age modifies the contribution of Nrg1 to recovery.

Taken together, these experiments allowed us to dissect the role of Nrg1 signaling at the cellular, circuit, and behavioral levels, and to establish its contribution to cortical repair and functional recovery after traumatic brain injury.

## Materials and methods

### Animals

For *in vitro* studies, we used CD1 (wild-type phenotype) and C57BL/6J Nrg1^tm3Cbm^floxed (named Nrg1 floxed) from Birchmeier lab (http://www.informatics.jax.org/allele/MGI:2447761) mouse embryos at E14.5 post-coitum for gain of function approaches. To study the role of Nrg1 signaling upon CCD, we used young (3-4-month-old) and mature (9-13-month-old) female mice from tamoxifen inducible transgenic mice to induce a ubiquitous deletion of Nrg1 (Nrg1-UBC-CreERT2) for loss-of function approaches. The mutation in Nrg1 gene is found in the exon 7, 8, and 9 of the EGF C-terminal, deleting all Nrg1 isoforms (Meyer & Birchmeier, 1995; Yang et al., 2001). All the animal experiments were supervised by the bioethical committee of Centro de Investigación Príncipe Felipe (Valencia, Spain), according to the European and Spanish bioethical regulations in terms of Animal Welfare (Ethical approval from the Conselleria of the Comunidad Valenciana register GVRTE-2019-641478, GVRTE-2020-1030394, GVRTE20212013496). The animals were group-housed with food and water ad libitum in standard housing conditions.

### Primary neuronal culture and neuron transfection by electroporation

Primary cortical neuron cultures were prepared from CD1 embryonic mice (E14.5-15.5), as previously described (Fazzari et al., 2010; Fazzari et al., 2014; Rodríguez-Prieto et al., 2021). Briefly, embryonic cortices were dissected and placed into ice-cold sterilized HBSS (7mM HEPES and 0.45% glucose). The tissue was disaggregated in trypsin-EDTA buffer at 37°C for 12 minutes. Cortices were washed with HBSS and gently homogenized for maximum 8-12 repetitions in 1.5ml of plating medium (Minimum Essential Medium (MEM) supplemented with 10% horse serum (Invitrogen), 0.6% glucose and antibiotics (penicillin, 10000 U/ml; streptomycin, 10 mg/ml)). Cell solution was filtered through a cell strainer (70µm). Cells were counted using a Neubauer chamber and subsequently seeded into 12-well plates pre-coated with poly-L-lysine (0.1 mg/mL, Sigma-Aldrich) at a density of 110,000 cells per well. Neuronal electroporation was performed with the NEPA21 (Nepa gene) system, as previously described (Rodriguez-Prieto et al, 2021). Briefly, an adequate volume of the cell suspension was centrifuged at 250g for 5 minutes in electroporation medium (Gibco Opti-MEM, 11524456, ThermoFisher Scientific). To obtain a concentration of 10^6^ cells/ml in each electroporation cuvettes, the pellet was resuspended in electroporation medium and mixed with the desired amount of DNA: pAAV-hSyn-RFP (3 μg, 22907, Addgene), pCMV-GFP-IRES-Cre (3 μg, Fazzari et al., 2010), CRD-Nrg1_FL and Nrg1_ICD (24 μg, Fazzari et al., 2014). The cuvettes were inserted in the electroporator under the pre-established conditions: poring pulse of 2 ms, 175 V, 50 interval, 10 decay rate, positive polarity, two times; transfer pulse of 50 ms, 20 V, 50 interval, 40 decay rate, positive and negative polarity, five times (Rodríguez-Prieto et al., 2021). After the electroporation, 45000-50000 cells per treated group were plated in 12-well plates. To reduce variability, we combined both control and treated groups in the same well. Control cells were electroporated with pCMV-GFP-IRES-Cre (GFP) and pAAV-hSyn-RFP (RFP) plasmids, while treated cells were electroporated with GFP and Nrg1-ICD or Nrg1-FL, according to each condition. Of note, the plasmids were obtained from transformed bacteria using a commercial kit (MB051, NZYMaxiprep). CRD-Nrg1 full length tagged with GFP was kindly provided by Prof. Bao Jianxin and cloned into pcDNA3.1 TOPO (Invitrogen). Nrg1-ICD was subcloned from pcDNA3.1 TOPO (Invitrogen) to pCAGEN (11160; Addgene) (Bao et al., 2003; Fazzari et al., 2014). Neuronal cultures were stored in a humidified incubator containing 95% air and 5% CO_2_. The plating medium was replaced 3 hours later with equilibrated neurobasal medium supplemented with B27, GlutaMAX (Gibco; Life Technologies Co.) and antibiotics (penicillin, 10000 U/ml; streptomycin, 10 mg/ml). To maintain the cells for longer studies, an additional ml of neurobasal medium was added to each well one week later and the medium was refreshed every two days until 14 days *in vitro*.

### Experimental *in vitro* model of axonal injury and quantification of axonal outgrowth

To study axonal regeneration upon injury *in vitro*, a protocol of physical injury *in vitro* was developed. Briefly, to reduce variability from injury protocol, a sparse labeling of the neurons was performed and intrinsic control was plated in each well. 4 days *in vitro*, physical insult was provoked by scratching the middle line of the coverslip using a mechanical pencil (0.5mm) in sterilized conditions. 72 hours and 10 days after the scratch, the cells were fixed and immunolabeled to strengthen the fluorescence. Pictures were taken with a Zeiss Axio Observer Z1 microscope, equipped with a Zeiss AxioCam MRm camera and an illumination system (Colibri), consisting on different excitation LED’s (385 for DAPI, 475 for 488, 567 for 555 and 630 for 647 fluorophores). The pictures were obtained using 20x NA 0.5 dry objective. Images were analyzed using a sholl analysis, by counting the number of axons crossing the empty space (10 days post-scratch: axons counted at 160, 240 and 320µm far from the border). These raw data were normalized using the following criterion. First, the number of axons crossing each line was transformed to axons/mm. Second, the data was normalized by the number of neurons counted per condition in different fields along the coverslip (Al-Ali et al., 2017). Lastly, this value was normalized to the quantification obtained in the control condition in the proximal line, which was assumed to be the baseline for axonal outgrowth.

### Experimental *In Vivo* Model of CCD

As previously described (González-Manteiga et al., 2022), 100µl of morphine (1.5mg/ml) was administered to mice 30 minutes before the surgery and 100µl of buprenorphine (0.03mg/ml) 6h later and the next morning. Moreover, weight loss was monitored along the experiment as an early humane endpoint parameter. Thus, animals that presented a body weight loss above 20% would be sacrificed and excluded from the study. Stereotactic injection of AAVs for the expression of GFP was performed to trace axonal projection of the corpus callosum as previously described (Navarro-González et al., 2019). Briefly, mice were placed in a stereotactic frame, under isoflurane anesthesia with the skull exposed. To target the primary motor cortex, the selected coordinates (mm) relative to the bregma were as follows: anteroposterior, 0.2; mediolateral, 1.5; and dorsoventral, −0.5. Then, the Hamilton syringe containing the virus was introduced 1mm into the cortex to produce a pocket. Two minutes later, the syringe was risen 0.5mm and 1μL of the virus (diluted 1:4) was injected at a flow rate of 0.1μl/min. After the injection, the needle was left up to 3 minutes for an appropriate virus diffusion. To induce Nrg1 deletion, a single dose of tamoxifen (8 mg; T5648-1G, Merck Life Science, S.L.) dissolved in corn oil (C8267-500ML, Merck Life Science, S.L.) was administered intraperitoneally to UBC-CreERT2; Nrg1^flox/flox^ mice seven days before injury induction. This protocol effectively triggers recombination and abrogates Nrg1 expression independently of genotype. In the cohort of mature female mice, half of the animals received tamoxifen (genotype: Cre-; Nrg1^flox/flox^), whereas the other half were treated with corn oil only (genotype: UBC-CreERT2/wt; Nrg1^flox/flox^), to maintain a sufficient number of control animals within the same experimental batch.

To perform the controlled cortical damage (CCD), a 2.1mm diameter drill tip was introduced in the right primary motor cortex at a constant speed of 0.5mm/s, using a motorized stereotactic instrument to reduce injury variation (Stoelting, 51730M). Considering the size of the drill, the coordinates relative to bregma (mm) suffer a slight antero-posterior modification, compared to the ones used for AAV injection: anteroposterior, 0; mediolateral, −1.5; and dorsoventral, −2.5. The drill was introduced into the brain 3 times, waiting 2min with the drill inside after the first immersion.

### Nrg1 construct and adenoassociated viral vectors

To study the role of Nrg1 signaling upon CCD *in vivo*, we labeled cortico-cortical projections using an adeno-associated viral vectors (AAV). The virus construct, production and validation were previously described (Fazzari et al., 2014; Navarro-González et al., 2019). The AAV vector was produced to express GFP under human Synapsin promoter. This vector for GFP expression was deposited by Bryan Roth into the Addgene repository (“Addgene: pAAV-hSyn-EGFP, reference number 50465”).

### Immunohistochemistry

Eight days after CCD, mouse brains were processed as previously described (González-Manteiga et al., 2022; Navarro-González et al., 2019). Briefly, mice were transcardially perfused with 4% PFA to collect the brain, following a 2h postfixation and cryoprotected in 30% sucrose for 72h. The brains were sectioned with a cryostat at 40μm (Cryostat Microm HM550, ThermoFisher Scientific), distributing the slices in 8 different anteroposterior ordered series for 3D reconstruction analysis. Primary and secondary antibodies used were diluted in PBS with 0.25% Triton and 4% BSA. Incubation with the primary lasted for 48h, while the secondary antibodies were left overnight at 4°C in agitation conditions. The antibodies and dilution used were as follows: anti-GFP (1:1000; GFP-1010, Ave Labs, Davis, CA, USA), anti-IBA1 (1:500, 234004, Synaptic Systems, Göttingen, Germany), anti-WFA biotinylated (1:1000, L1516-2MG, Sigma Aldrich, St Louis, MI, USA), anti-chicken 488 (A-11039, Thermofisher, Waltham, MA, USA), anti-guinea pig 647 (A21450, Invitrogen), and streptavidin CyTM5 (PA45001, GE Healthcare, Chicago, IL, USA). . All the secondary antibodies were diluted 1:500. Fluorescence imaging of the whole slides was performed using a Leica Aperio Versa at 10× NA 0.32 Plan Apo, equipped with a camera model Andor Zyla.

### Bioinformatic image processing

Image stacks were processed using a semi-automated workflow previously described (González-Manteiga et al., 2022). Briefly, fluorescence images acquired with a Leica Aperio Versa scanner were separated by channel and processed in Fiji to generate binary DAPI masks defining the slice contour. The DAPI channel was smoothed with a Gaussian blur (σ = 3) and thresholded to obtain full-slice masks that excluded tissue holes caused by CCD injury. Masks were applied to all channels to normalize field dimensions, and slice areas were automatically profiled and sorted using custom Python scripts.

Slices spanning 2.5 mm to –1 mm from bregma (Paxinos and Franklin atlas) were reconstructed in 3D and aligned along the midline using an adapted “Align Image by Line” Fiji plugin. The alignment step was automated by a custom macro, which achieved approximately 95% accuracy; manual verification was performed in cases of tissue distortion or incomplete slices.

### Axonal preservation analysis

GFP-immunostained coronal sections were analyzed using ImageJ to quantify fluorescence intensity along the cortical layers adjacent to the lesion border. Only samples with clear viral labeling at AP 0.2 and LM 1.5 (Paxinos and Franklin atlas) were included; sections with off-target or weak labeling were excluded. Comparable slices between AP +0.86 and 0 mm were selected from the right primary motor cortex near the CCD lesion, and equivalent homotopic regions were analyzed in uninjured controls for comparison.

To evaluate axonal preservation, horizontal regions of interest (ROIs) were placed parallel to the injury border to measure GFP intensity across cortical layers II/III, IV, and V. The preservation ratio was calculated as the mean signal intensity in pixels near versus far from the lesion (close/far ratio). Data were normalized to total ROI intensity to minimize inter-slice variability and analyzed using two-way ANOVA.

This quantitative approach provided a layer-resolved measure of axonal loss and remodeling following CCD in control and Nrg1-deficient mice.

### Stereological measurements and image analysis

From sorted and aligned stacks, the quantification of injury volume, inflammation, and tissue damage impact was done. The methodology used was as previously described (Navarro-Gonzalez et al., 2021). Briefly, the injury area was measured manually and the volume of the injury was lately calculated by the integration of its polynomial fit equation, according to the formula: V = ∫ (𝑥)x in which A is the cross-sectional area and x represents the width of the interval [a,b]. To estimate inflammation, IBA1 staining pictures were used. Briefly, Gaussian blur filter (sigma = 5) was set to facilitate the homogenization and detection of inflamed area. After that, a threshold was manually set, selecting the value that better fit in the whole samples, according to the contralateral hemisphere. The image was converted into a binary mask in which the region of interest per slice was automatically identified using the function named “Analyze particles” (size = 1000-infinity). The area was calculated on the region of interest detected by this tool. From these values, the volume of each parameter was calculated by integration of the polynomial fit equation, as previously described.

To analyze the cell-to-cell interaction between microglia and PNN upon CCD, the number and intensity of WFA+ cells was analyzed in the PL, determined by IBA1+ labeling. Briefly, rectangles in both IBA1+ and IBA1− (somatosensory cortex) were selected as regions of interest (ROI) in the ipsilateral and contralateral cortex. To delimit the number of WFA+ cells, the same threshold was established on both ROIs in the IBA1+ or IBA1− area, according to contralateral staining which becomes the intrinsic control in each sample. Once the threshold was selected, we automatically detected the contour of the WFA+ cells, using the tool “Analyze particles” (size = 200-infinity) of ImageJ. The ROIs of each cell were saved, and the intensity was measured in the raw images. Data from ipsilesional regions was normalized to the respective contralateral ROIs for each sample for CCD characterization and LOF ipsilesional regions, while data was normalized to WT group in the contralesional areas of LOF approaches. Data represent the fold change in the graph (mean ± SEM).

### IBA1 immunofluorescence imaging and analysis

To further assess the role of endogenous Nrg1 in neuroinflammation, we implemented an additional fluorescence-based approach to quantify microglia activation through IBA1. Brain slices were imaged using a Zeiss ApoTome.2 system equipped with a 20× objective (AxioCam MRm, NA: 0.35). Images were acquired in full z-stack mode across fluorescence channels (AF488 475 nm, CY5 630 nm, DAPI 385 nm,).

For each slice, images were acquired from both the lateral and central regions relative to the lesion site. Using Fiji software, two rectangular (approx. 100x400-500 μm) regions of interest (ROIs) were selected at 500 µm from the lesion, respectively, to assess potential differences in inflammation levels at varying distances from the injury. Each ROI consisted of a 16-bit z-stack image comprising the fluorescent channels. The IBA1 channel was isolated, and a common minimum/maximum intensity range was applied across all images using the brightness/contrast settings. The images were then converted to 8-bit format, and the Fire lookup table (LUT) was applied to enhance cell visualization.

Two different analyses were performed. First, a custom macro was developed in ImageJ to apply incrementally increasing intensity thresholds (from 20 to 240, in steps of 10). For each threshold level, the mean pixel intensity was determined. Since no statistical difference was found between lateral and central images, the results were pooled together. Importantly, threshold values exceeding 100 were not considered, as they led to significant loss of signal.

Secondly, the threshold level that best isolated the cell soma was manually selected for each image. Using the “Analyze Particles” function (size = 30–infinity), five morphological and intensity parameters were measured for each cell: perimeter and integrated density. ROIs that did not correspond to individual soma or that were located at the image borders were manually excluded. The integrated density values were normalized to the average value of the control animals. Since no statistical difference was found between lateral and central images, the results were pooled together.

### Motor behavior analysis

#### Wide-spaced ladder test

Motor coordination and skilled locomotion were assessed using a wide-spaced ladder test as described (González-Manteiga et al., 2022). The apparatus consisted of an enclosed transparent walkway (MotoRater system, TSE Systems) equipped with two mirrors positioned above to enable simultaneous observation from three angles, and a high-resolution camera placed below for video acquisition. Mice were allowed to traverse a horizontal ladder with rungs spaced 3 cm apart, performing three consecutive runs per session. Videos were analyzed using a 0–5 scoring scale based on the number of paw placement errors and gait symmetry. Gait symmetry was defined as the alignment of the hindlimb step with the position previously occupied by the forelimb.

#### Rotating pole test

Motor performance following cortical injury was evaluated using the rotating pole test, a sensitive assay for detecting unilateral motor deficits, as described (Talhada et al., 2019; González-Manteiga et al., 2022). Mice were trained to traverse a wooden pole elevated 1 m above the table surface under three conditions: static (no rotation), and rotation to the right or to the left at 3 rpm. Tests were always conducted in the same order, from static to right and finally left rotation, the latter being the most challenging given that the CCD was performed on the right motor cortex (thus affecting the contralateral, left hindlimb). Performance was video recorded and scored on a 0–6 scale according to walking ability and the number of paw slips, as previously described (Talhada et al., 2019; González-Manteiga et al., 2022).

### Statistical analysis

Statistical analysis and graph preparations of all the figures were done using Graph Pad Prism 9 software. The data is represented as mean ± SEM. The significance is indicated with an asterisk in each figure legend, considering p ≤ 0.05 as significant. The statistical test used for each analysis is also mentioned in each figure legend. Normality distribution of the samples was assessed using D’Agostino-Pearson and Shapiro-Wilk test to select the most accurate statistical test. For the quantification of axonal regeneration *in vitro*, a two-way ANOVA with repeated measurements and Sidàk’s multiple comparison tests were used to compare between the different groups. All the data was normalized to the number of electroporated cells per group and to control condition. The behavioral data analysis was assessed using a two-way ANOVA with repeated measurements and Sídák’s multiple comparison tests to compare Ctrl and Nrg1 KO within different time points in both RPT and WSL. Noteworthy, the data was normalized to the pre-injury data for the column bar graphs in the RPT test. To analyze the Nrg1 role in injury impact and neuroinflammatory response *in vivo*, we used an unpaired t-test to compare the IBA1^+^ area and cell morphology and intensity values between groups, while a 2 way ANOVA was used to compare the mean intensity values for each threshold between groups. To analyze perineuronal nets upon CCD in the LOF studies, we calculated the fold change of WFA+ cell number decrease in the ipsilesional area relative to the contralesional area within each slice and compared both phenotypes using Mann-Whitney U test. For the contralesional region, we compared both WFA+ cell number between groups using a Mann-Whitney U test. Finally, to assess signal intensity measurements, two-way ANOVA with repeated measurements and Sídák’s multiple comparison tests were performed to compare cortical profile distribution between groups and fold change of intensity increase for axonal preservation within layers.

### Linear Discriminant Analysis

Multivariate relationships among injury and inflammatory features were assessed using Linear Discriminant Analysis (LDA), implemented in Python (version 3.11) with the scikit-learn library (version 1.7.2). For the analysis we used the classical svd solver, corresponding to Fisher’s Linear Discriminant Analysis, which identifies the linear combination of features that maximizes between-group variance relative to within-group variance.

For each animal, an LDA score was computed to represent its position along the discriminant axis separating CTRL and Nrg1 KO genotypes. Group separation along LDA score was statistically evaluated using a two-tailed t-test, and model performance was validated by leave-one-out cross-validation.

The Python code used for these analyses was written with the assistance of ChatGPT (OpenAI, 2025) to streamline implementation. All computational steps, parameters, and results were independently verified and rechecked in full by the authors.

## Results

### Nrg1 cell-autonomously promotes axonal outgrowth upon injury *in vitro*

Neuregulin-1 (Nrg1) plays a critical role in neuronal development, but its potential in axonal regeneration remains poorly understood. Building upon our previous findings demonstrating Nrg1’s importance in cortical axonal growth (Rodríguez-Prieto et al., 2024), we investigated whether Nrg1 can directly promote axonal regeneration following injury.

To examine the cell-autonomous role of Nrg1 in axonal regeneration, we employed a gain-of-function (GOF) approach. This involved expressing full-length Nrg1 (Nrg1-FL) or its intracellular domain (Nrg1-ICD) in primary neurons. We and others have previously shown that Nrg1-ICD expression effectively activates Nrg1 intracellular signaling (Bao et al., 2003; Fazzari et al., 2014; Navarro-González et al., 2019; Rodríguez-Prieto et al., 2024).

For single-cell resolution and analysis of neuronal response to injury, we co-cultured electroporated neurons expressing Nrg1-FL or Nrg1-ICD (GFP-labeled) with non-electroporated control cells (Ctrl, RFP-labeled), as previously described (Rodríguez-Prieto et al., 2024, Rodríguez-Prieto et al., 2021). This method enabled the evaluation of the cell-autonomous effects of Nrg1 and its intracellular signaling (Figure 1A,B). Briefly, we induced scratch injury at day 4 *in vitro* (DIV4) and analyzed axonal outgrowth 10 days later (DIV14) (Figure 1B). Namely, we quantified the number of axons crossing the lesion zone at proximal (P, 160 μm), medial (M, 240 μm), and distal (D, 320 μm) distances from the injury site (Figure 1C).

**Fig. 1.**
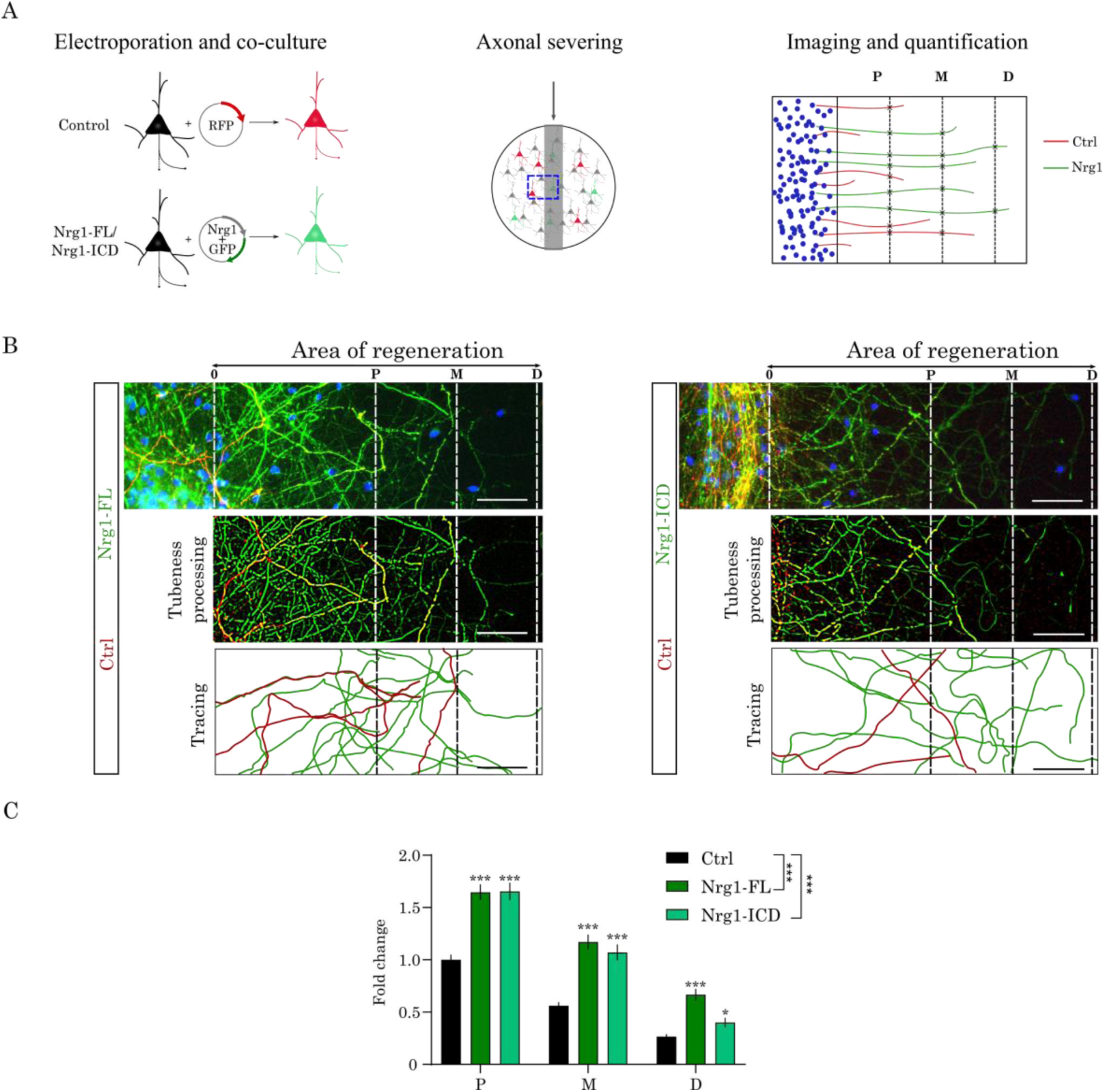
Nrg1 promotes axonal outgrowth *in vitro* after axonal injury. (A) Experimental design: primary cortical neurons were electroporated at DIV0 with GFP and RFP (control) or GFP together with Nrg1-FL or Nrg1-ICD plasmids. At DIV4, axons were mechanically severed, and regeneration was analyzed at DIV10. Axonal crossing was quantified at proximal (P, 160 µm), medial (M, 240 µm), and distal (D, 320 µm) distances from the lesion, using RFP+ axons as internal control. (B) Representative images of axonal regeneration in control, Nrg1-FL, and Nrg1-ICD neurons, with manual tracing of regenerated axons (scale bar = 50 µm). (C) Quantification of axonal outgrowth shows significantly increased axonal crossing in Nrg1-FL and Nrg1-ICD neurons compared with controls at all distances. Bars represent mean ± SEM (Ctrl, n = 270 axons; Nrg1-FL, n = 90; Nrg1-ICD, n = 90; 3 independent experiments). Statistical analysis: two-way repeated measures ANOVA with Sidák’s post hoc test (*p ≤ 0.05, ***p ≤ 0.001).

Quantification of the injured area revealed a robust increase in Nrg1-FL and Nrg1-ICD axon regeneration compared to the control group. Notably, a substantial increase in axonal growth was consistently observed in Nrg1-FL and Nrg1-ICD expressing neurons at all measured distances (P, M, and D) from the lesion site (Figure 1B,C). These results demonstrate that Nrg1, particularly the activation of its intracellular signaling, is sufficient to promote axonal outgrowth following injury *in vitro*.

These observations provided a compelling proof of principle, suggesting that Nrg1 intracellular signaling may play a significant role in the axonal response to injury, and prompted us to follow up this investigation *in vivo*.

### Nrg1 deficiency exacerbates cortical wiring deficits after CCD *in vivo*

Next, we investigated the role of Nrg1 in cortical injury, a context in which the endogenous Nrg1/Erbb4 signaling pathway remains unexplored. Early developmental deletion of either Nrg1 or its receptor Erbb4 induces various brain developmental alterations (Mei and Nave, 2014), which complicate interpreting Nrg1’s role in the adult brain. To circumvent this limitation, we developed a conditional mouse model by crossing *Nrg1*^flox/flox^ mice with UBC-CreERT2 mice (UBC-CreERT2; *Nrg1*^flox/flox^; hereafter Nrg1 KO) to enable adult-specific Nrg1 deletion (Figure 2A). Quantitative PCR analysis confirmed efficient recombination, showing a dramatic reduction in *Nrg1* mRNA expression following tamoxifen administration (Ctrl: 1.00 ± 0.24; Nrg1 KO: 0.047 ± 0.016 a.u.; mean ± SEM; n = 3 per group; p = 0.014, unpaired t-test).

**Fig. 2.**
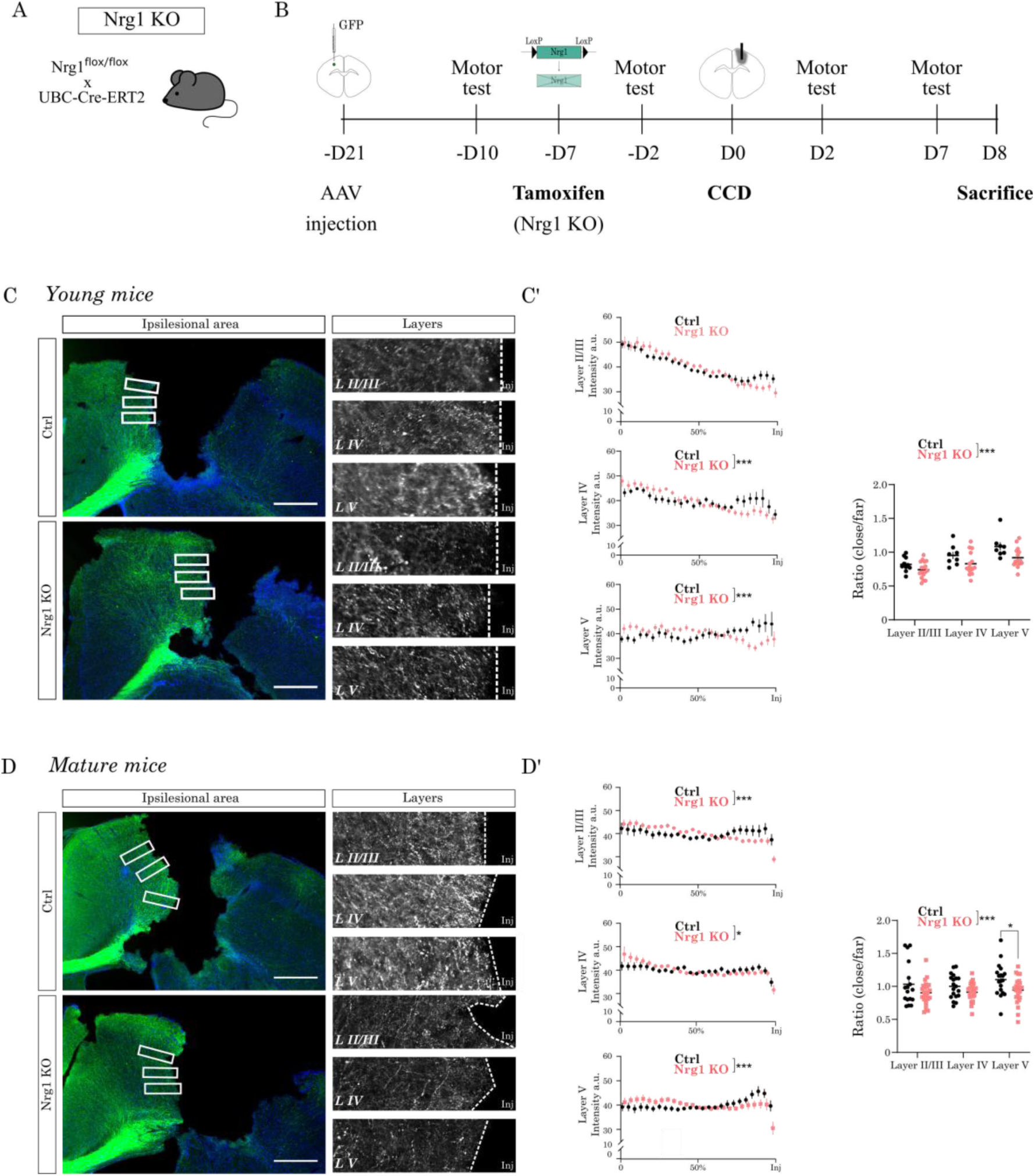
Nrg1 deficiency exacerbates cortical wiring deficits after CCD *in vivo*. (A, B) Experimental design: callosal projections were labeled by injecting AAV-Syn::GFP into the left motor cortex. Then, tamoxifen was administered to induce acute Nrg1 deletion prior to controlled cortical damage (CCD) in in UBC-CreERT2;Nrg1flox/flox (Nrg1 KO) mice. After controlled cortical damage (CCD) in the right motor cortex, mice underwent motor testing and were subsequently analyzed histologically. (C) Representative images of GFP+ callosal axons (green) in the perilesional (PL) region of young mice, with boxed areas magnified on the right. DAPI (blue) was used as counterstain. Scale bar = 500 µm. (C′) Quantification of axonal density profiles close to the injury across cortical layers shows reduced preservation in Nrg1 KO compared to controls. Ratios of axonal density near versus distant from the lesion confirm a consistent loss of callosal projections in Nrg1-deficient young mice. Bars represent mean ± SEM (Ctrl, n = 10 sections; Nrg1 KO, n = 17 sections). Statistical analysis: two-way ANOVA with Sidák’s multiple comparisons test (***p ≤ 0.001). (D) Representative images of GFP+ callosal axons in mature mice, with boxed areas magnified on the right. Scale bar = 500 µm. (D′) Quantification as in (C′) shows significantly reduced axonal density in Nrg1 KO mice compared to controls. Bars represent mean ± SEM (Ctrl, n = 17 sections; Nrg1 KO, n = 25 sections). Statistical analysis: two-way ANOVA with Sidák’s multiple comparisons test (*p ≤ 0.05, ***p ≤ 0.001).

We employed a refined controlled cortical damage (CCD) model previously described (González-Manteiga et al., 2022). The CCD model is based on a mechanical injury that in our experience results in a very consistent and reproducible cortical injury. We examined injury responses in both young (3-4 months) and mature (10-12 months) mice to comprehensively understand Nrg1’s role in cortical damage across different ages.

As a model to study axonal remodeling after injury, we focused on callosal connections (CC). CC are the main interhemispheric projections and they are critical for motor coordination (De León Reyes et al., 2020; Empl et al., 2022). To trace CC, we utilized viral vectors expressing GFP under the human synapsin promoter, specifically labeling neurons in the left primary motor cortex and their contralateral callosal projections (Figure 2B). We previously showed that acute Nrg1 deletion in these mice does not alter the baseline contralateral callosal projection profile (Rodríguez-Prieto et al., 2024).

We performed CCD injury in the right motor cortex using a stereotactic apparatus (Figure 2B). We found that the volume of the CCD injury was similar in control and Nrg1 KO mice both in young and mature mice (Supplementary Fig. S1B).

Given the increased vulnerability of axons near the injury border, we analyzed axonal density at varying distances from the lesion to clarify Nrg1’s role in injury response. Our analysis revealed that Nrg1 loss consistently resulted in decreased callosal axon density near the injury site (Figure 2C-D’). The axonal density ratio near the lesion was significantly lower in Nrg1 KO mice compared to controls, this finding was robustly reproduced across both young and mature experimental groups (Figure 2C-D’). Reduced axonal density in Nrg1-deficient mice indicates a failure to preserve or remodel callosal connections, potentially impairing motor coordination after injury.

In summary, acute Nrg1 deletion significantly affected contralateral cortical wiring following cortical damage. Specifically, Nrg1 deficiency resulted in a pronounced decrease in axonal density proximal to the injury site.

### Inflammatory response to CCD in Nrg1-deficient mice

Our experiments indicated a role for Nrg1 signaling in axonal growth and in the response of cortical circuits after CCD. However, previous studies have also suggested a potential involvement of Nrg1 in neuroinflammation after stroke. Notably, most of these studies focused on the administration of soluble exogenous Nrg1, and did not address the role of endogenous Nrg1 and its intracellular signaling. Additionally, the anti-inflammatory properties of Nrg1 have been primarily investigated in models of ischemic and traumatic brain injury (MCAO and experimental TBI) (Guan et al., 2015; Noll et al., 2019; Xu et al., 2005, 2004; Zhang et al., 2018), where the extent of brain damage is considerably larger than in CCD. Hence, here we investigated the inflammatory response in Nrg1 deficient mice upon CCD. To this end, we used IBA1 immunostaining as a marker for microglia/macrophage activation.

We assessed inflammation in both young (3–4 months) and mature (10–12 months) mice to determine whether the role of Nrg1 varies with age. To study the activation of the microglia, we first evaluated the overall IBA1+ volume in the perilesional region. The volume and intensity of IBA1+ staining showed a tendency to increase in Nrg1 deficient mice in both young and mature mice. To gain further insights, we performed a multivariate examination of these data by linear discriminant analysis (LDA) considering the volume of the injury, the volume IBA1+ area and integrated density of IBA labeling. LDA indicated a genotype-specific separation between control and Nrg1 KO mice that appeared more significant for the group of mature mice (Supplementary Figure 1A-H).

Next, we performed a morphometric analysis of the soma of the microglia and the relative coverage of the IBA1 labeling. As expected, we observed a gradient in IBA1 staining intensity and macrophage/microglial cells recruitment, with stronger signals and higher cell densities near the lesion site, gradually decreasing in more distal regions. Hence, we focused our analysis on an intermediate zone of microglial activation located 500 µm from the lesion site. Specifically, we quantified the perimeter of the soma, the intensity of IBA expression in the soma, and the relative coverage of IBA+ area. Overall, we failed to observe differences in these parameters in young mice (Fig. 3A-C). Conversely, we found a significant increase in somatic IBA1 expression in mature mice, as measured by integrated density of the labeling, in Nrg1 deficient mice as compared to control (Fig. 3D-E). Finally, we quantified IBA1+ coverage by measuring the percentage of image area occupied by IBA1 signal across increasing intensity thresholds, which revealed higher values in Nrg1-deficient mice compared with controls (Fig. 3F).

**Fig. 3.**
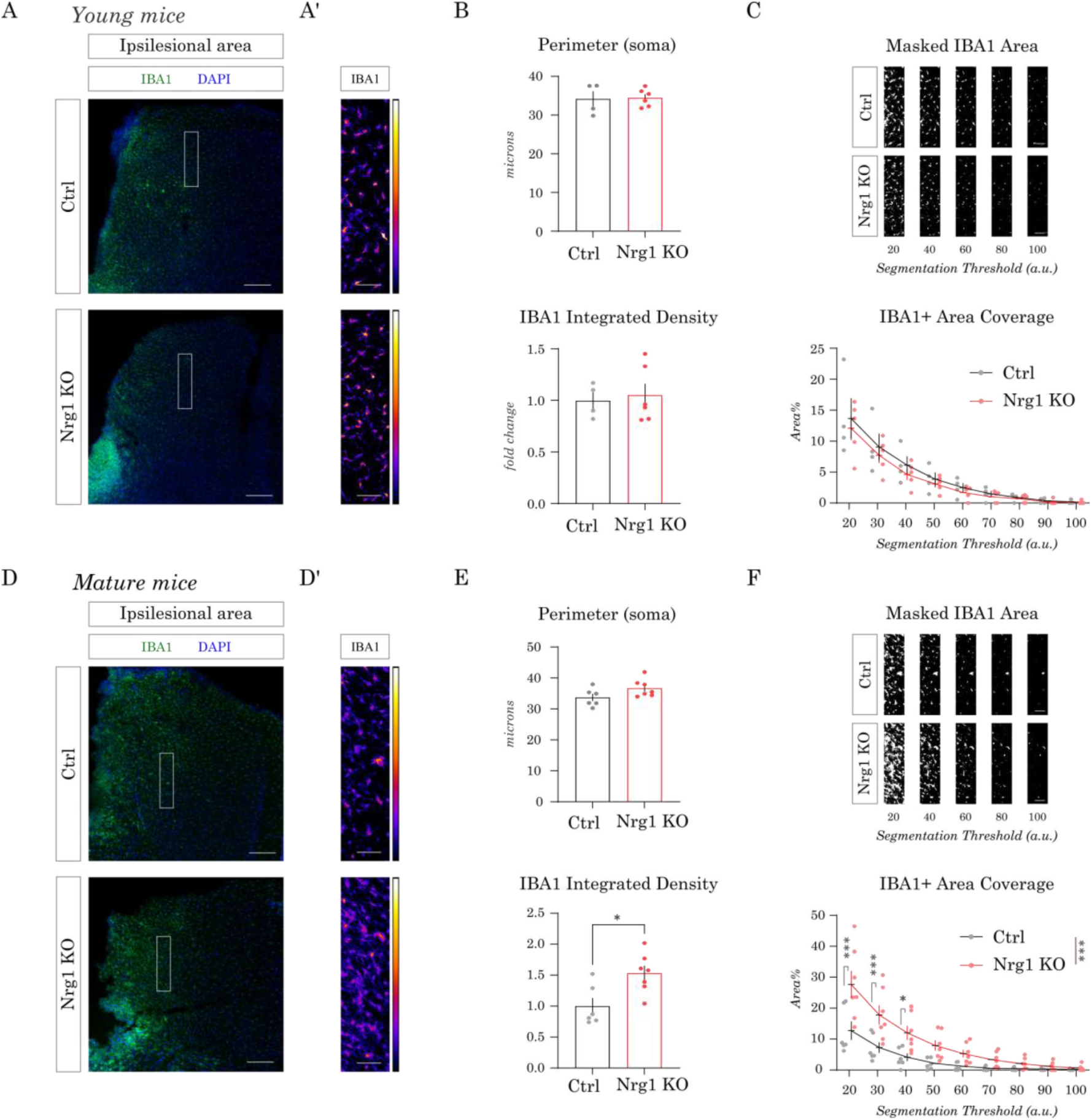
Inflammatory response to CCD in Nrg1-deficient mice. (A, A′) Young mice. Representative IBA1 immunostaining (gray) with DAPI counterstain (blue) in the perilesional cortex (A). The boxed region is shown in A′ as a color-coded image, with quantifications in (B, C). (A) Scale bar = 100 µm. (A’) Scale bar = 50 µm. (B, C) Quantification of IBA1 labeling in young mice. (B) Microglial soma perimeter and IBA1 integrated density. (C) Area coverage of IBA1⁺ staining. Top: segmented images at increasing intensity thresholds. Bottom: quantification of IBA1⁺ area coverage expressed as percentage of total area. Bars represent mean ± SEM (Ctrl, n = 4 mice; Nrg1 KO, n = 6 mice). Statistical analysis: unpaired Student’s t-test for soma perimeter and integrated density; two-way repeated measures ANOVA with Sidák’s post hoc test for area coverage. (D, D′) Mature mice. Representative IBA1 immunostaining (gray) with DAPI (blue) in the perilesional cortex (D). The boxed region is shown in D′ as a color-coded image, with quantifications in (E, F). (D) Scale bar = 100 µm. (D’) Scale bar = 50 µm. (E, F) Quantification of IBA1 labeling in mature mice. (E) Microglial soma perimeter and IBA1 integrated density, showing a significant increase in integrated density in Nrg1 KO mice compared with controls. (F) Area coverage of IBA1⁺ staining. Top: segmented images at increasing thresholds. Bottom: quantification of IBA1⁺ area coverage (%). Bars represent mean ± SEM (Ctrl, n = 6 mice; Nrg1 KO, n = 7 mice). Statistical analysis: unpaired Student’s t-test for soma perimeter and integrated density; two-way repeated measures ANOVA with Sidák’s post hoc test for area coverage (*p < 0.05, ***p < 0.001).

These findings correlate with previous data pointing out that elderly population are more susceptible to inflammatory response upon injury and, hence, poor functional outcomes (Shen, et al. 2019; Wangler et al. 2022).

Overall, these results indicate that Nrg1 modulates the inflammatory response after CCD, with mature mice showing greater dependence on its neuroprotective role than young mice.

### Nrg1 regulates PNN integrity after CCD

Perineuronal nets (PNNs) are specialized extracellular matrix structures composed of various proteoglycans, which contribute to the maturation and stabilization of synaptic circuits during CNS development (Devienne et al., 2021; Quattromani et al., 2018; Wang and Fawcett, 2012). Notably, PNNs preferentially enwrap fast-spiking parvalbumin (PV) interneurons, whose function is tightly regulated by Nrg1/ErbB4 signaling (del Pino et al., 2013; Fazzari et al., 2010; Mei and Nave, 2014; Navarro-Gonzalez et al., 2021). While the role of PNNs in the context of CNS injury remains unclear, previous studies, including ours, have reported a decrease in PNNs near sites of neuronal injury (Alia et al., 2016; González-Manteiga et al., 2022; Hsieh et al., 2017; Quattromani et al., 2018).

In the CCD model, this decrease in PNNs appeared particularly pronounced (González-Manteiga et al., 2022). Therefore, we investigated whether endogenous Nrg1 may also affect the degradation of PNNs after CCD. To address this question, we labeled PNNs with the specific marker Wisteria Floribunda Agglutinin (WFA) and quantified the number of WFA+ cells in both control and Nrg1 KO mice after CCD. We specifically analyzed both the PL region in ipsilesional side and the area in the contralesional hemisphere where contralateral projecting callosal neurons are located (Fig. 4A).

**Fig. 4.**
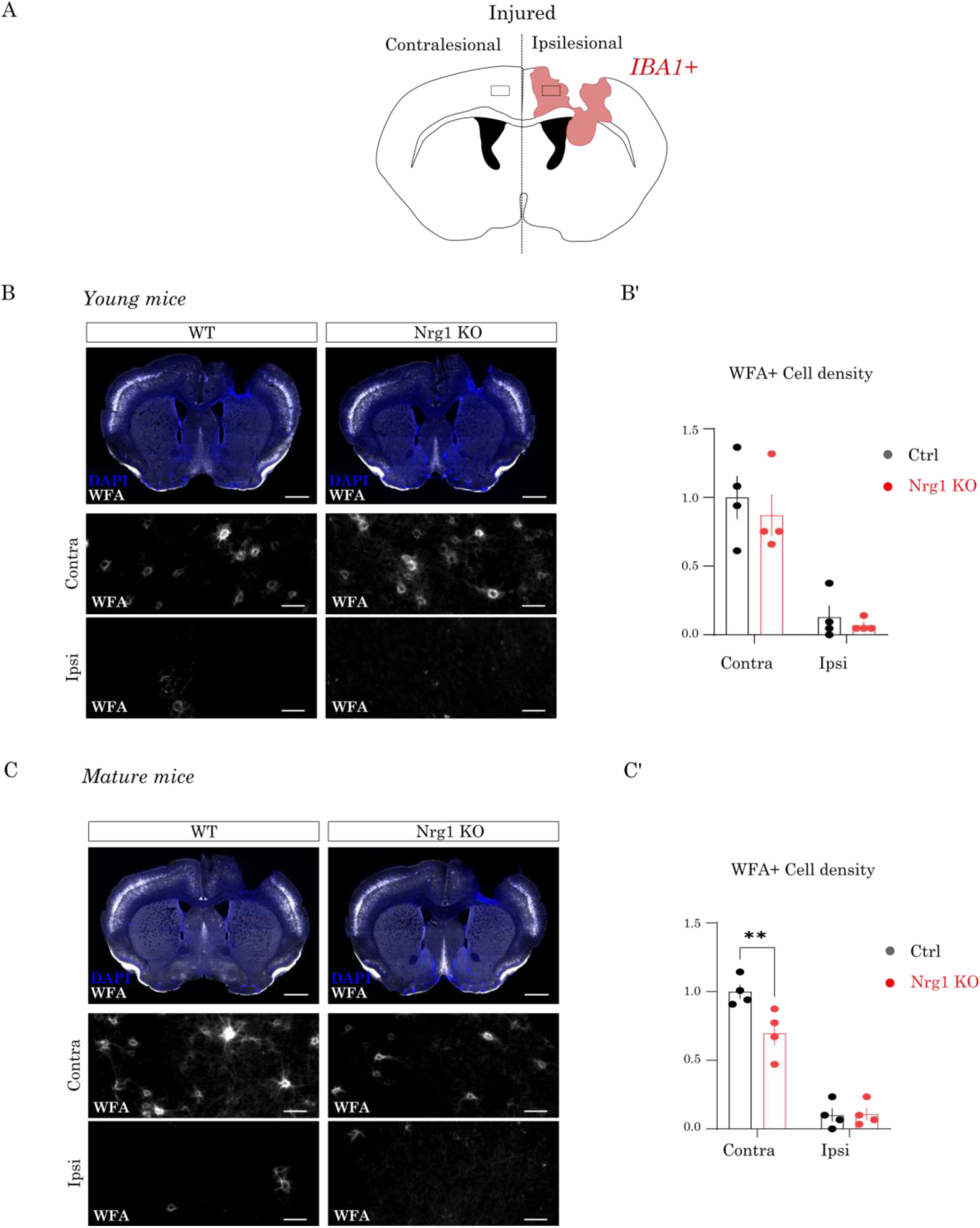
Nrg1 regulates perineuronal net (PNN) integrity after CCD. (A) Schematic illustrating the regions of interest (ROIs) selected for PNN quantification in ipsilesional and contralesional cortex. (B) Young mice. Top: representative WFA staining (white) in injured brains of WT and Nrg1 KO mice (scale bar = 1 mm). Bottom: higher-magnification images of ROIs (boxed in A) showing reduced WFA⁺ cells in the ipsilesional area (scale bar = 50 µm). (B′) Quantification of WFA⁺ cell number and fluorescence intensity in the contralesional area (left) and WFA⁺ cell number in the ipsilesional region (right). Data are expressed as fold change relative to control. Bars represent mean ± SEM (n = 4 animals per group). Statistical analysis: Two-way ANOVA. (C) Mature mice. Top: representative WFA staining in injured brains of WT and Nrg1 KO mice (scale bar = 1 mm). Bottom: higher-magnification ROIs (as in A) showing reduced WFA⁺ cells in the ipsilesional area (scale bar = 50 µm). (C′) Quantification of WFA⁺ cell number and intensity in the contralesional area (left) and WFA⁺ cell number in the ipsilesional region (right). Data are expressed as fold change relative to control. Bars represent mean ± SEM (n = 4 animals per group). Statistical analysis: Two-way ANOVA (**p < 0.05).

**Fig. 5.**
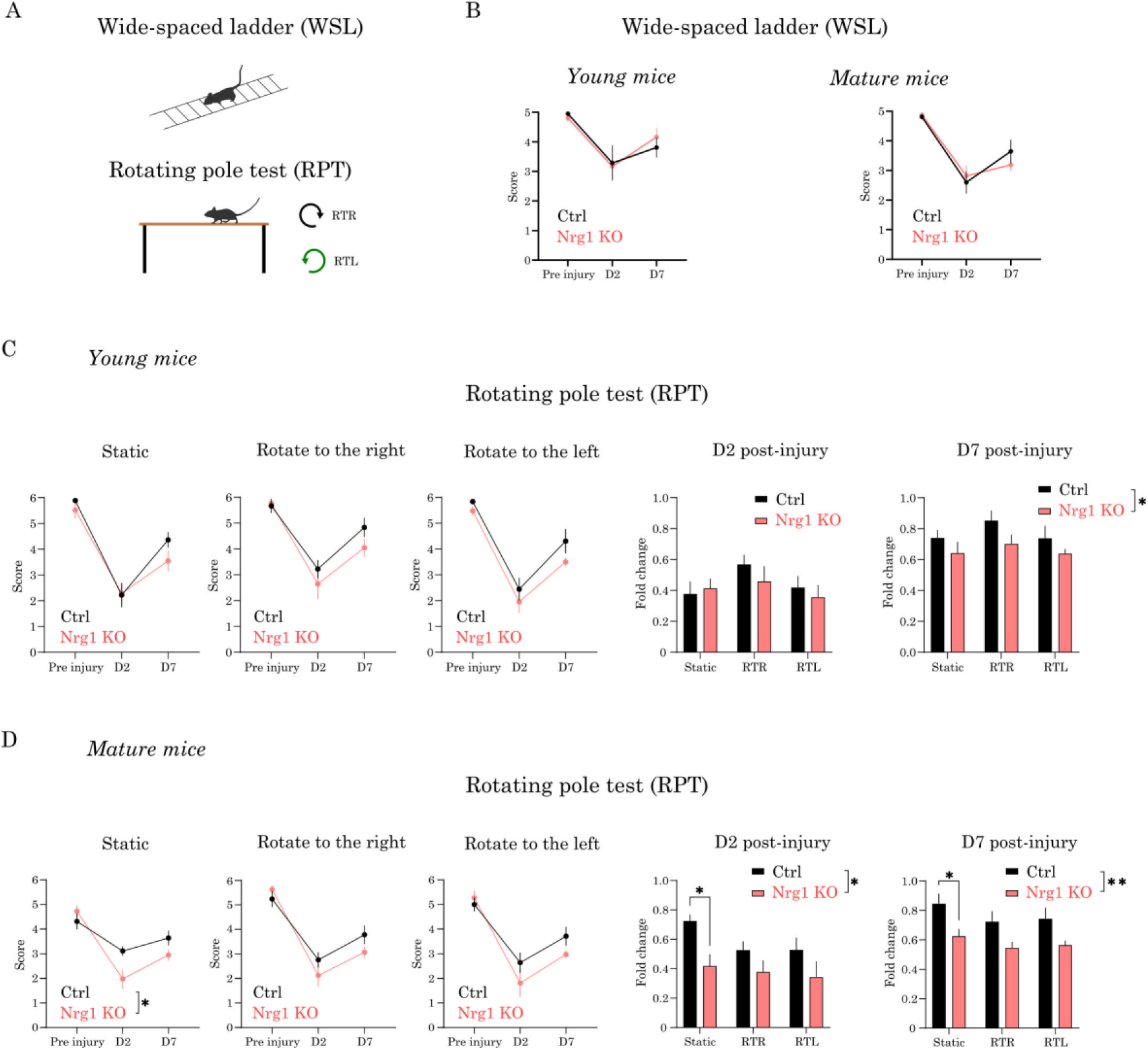
Nrg1 deficiency impairs motor recovery after CCD. (A) Schematic representation of the motor tasks used: wide-spaced ladder (WSL, horizontal) and rotating pole test (RPT). (B) WSL performance. Error rate was measured before injury (Pre), and at 2 days (D2) and 7 days (D7) post-injury in young (left) and mature (right) mice. CCD impaired performance in both genotypes, but no significant differences were detected between control and Nrg1 KO mice (C, D) RPT performance. Success rate was scored under static conditions and during rotation to the right (RTR) or to the left (RTL) before injury (Pre), and at D2 and D7 post-injury. Nrg1 KO mice performed significantly worse than controls in both young (C) and mature (D) groups, with more pronounced deficits in mature mice. Bars represent mean ± SEM (young: Ctrl, n = 6; Nrg1 KO, n = 7; mature: Ctrl, n = 6; Nrg1 KO, n = 7). Statistical analysis: two-way repeated measures ANOVA with Sidák’s post hoc test (*p < 0.05, **p < 0.01).

Consistent with our previous findings (González-Manteiga et al., 2022), we observed marked PNN degradation in the ipsilesional PL region, as evidenced by a major reduction in WFA+ cells in both young and mature mice. Specifically, seven days after CCD, WFA labeling was reduced by over 80% in the PL cortex of both age groups (Fig 4B’, C’). No significant differences in perilesional WFA staining were detected between control and Nrg1-deficient mice, possibly due to the dramatic decrease in WFA labeling observed in both genotypes.

In the contralesional hemisphere, mature Nrg1 deficient mice showed a significant decrease in WFA labeling density compared to control mice (Fig. 4C, C’). Additionally, we observed a non-significant reduction in the number of WFA+ cells. Similarly to what we observed for the microphage/microglial activation, Nrg1 deficient young mice did not show notable differences as compared to controls (Fig. 4B, B’).

In conclusion, these results confirm the substantial impact of CCD on PNN integrity and remodeling, which is often associated with plasticity. Furthermore, these findings suggest that endogenous Nrg1 may be involved in maintaining PNN integrity in an age-dependent manner, with more pronounced effects in mature mice.

### Nrg1 deficiency impaired motor skills after CCD

Taken together, our cellular and molecular analyses revealed alterations in neuronal circuitry in Nrg1-deficient mice after CCD. To determine whether these circuit alterations translate into functional deficits after CCD, we assessed motor recovery.

The CCD model involves a focal mechanical lesion targeting the region of the hindlimb motor cortex. We previously demonstrated that this injury impairs skilled motor tasks, such as the wide-spaced ladder (WSL) and the Rotating Pole Test (RPT) (González-Manteiga et al., 2022). The RPT is particularly demanding for motor coordination, requiring the mouse to walk forward while the pole is rotating either right (clockwise) or left (counterclockwise). This feature makes the RPT as an effective and reliable method to assess motor deficits caused by unilateral injuries, such as CCD.

To investigate whether Nrg1 promotes functional recovery upon brain damage, we next evaluated motor performance in Nrg1 deficient young and mature mice as compared to controls. Notably, the mice did not present any alterations in motor behavior before and after tamoxifen administration prior to injury induction (not shown). Therefore, the observed deficits are entirely attributable to impaired functional motor recovery following injury.

We evaluated motor behavior two days before the injury (-D2), and at D2 and D7 post-injury in young and mature female mice. Although we detected clear motor impairment in the WSL test after injury, we did not observe significant genotype-related differences. Conversely, we found that Nrg1 deficiency impaired motor performance in the RPT test in both young and mature mice. Consistent with the histological analysis of the cortical circuits, the differences between Nrg1 deficient mice and controls were more pronounced in mature mice compared to young animals, both at D2 and D7 after CCD.

Altogether, these impairments in motor function after CCD, coupled with our molecular and cellular analyses, demonstrate that endogenous Nrg1 plays a critical role in preserving neural circuit function and motor activity after CCD, with more pronounced effects in mature animals.

## Discussion

The present study elucidates the critical role of endogenous Neuregulin 1 (Nrg1) in cortical circuit response and functional recovery following traumatic brain injury. To our knowledge, this is the first study to use an inducible model of Nrg1 deficiency to specifically ablate Nrg1 expression in the adult brain, thereby avoiding the developmental confounds associated with constitutive loss-of-function models. Moreover, using a combination of *in vitro* and *in vivo* approaches, we demonstrate that Nrg1 signaling promotes axonal outgrowth in a cell-autonomous manner and modulates structural and functional outcomes after controlled cortical damage (CCD). Together, our data support the view that Nrg1 is required for maintaining axonal connectivity, preserving perineuronal net (PNN) integrity in mature mice, and sustaining motor function. These findings advance our understanding of Nrg1’s physiological role in post-injury plasticity and highlight its therapeutic potential for traumatic brain injury.

### Cell-autonomous role of Nrg1 in axonal response to injury

While canonical Nrg1/ErbB4 signaling has been extensively studied in development and schizophrenia (Mei and Nave, 2014; Mei and Xiong, 2008), our findings emphasize the underexplored role of non-canonical intracellular Nrg1 signaling in brain injury.

Prior studies have focused on exogenous Nrg1 administration (Deng et al., 2019; Gu et al., 2017); in contrast, our work demonstrates that endogenous Nrg1, acting through its intracellular domain (Nrg1-ICD), directly enhances axonal growth. The enhanced axonal outgrowth in neurons expressing either full-length Nrg1 or Nrg1-ICD supports the idea that intracellular Nrg1 signaling is sufficient to enhance intrinsic regenerative capacity.

This *in vitro* evidence prompted us to examine its relevance *in vivo*, where we found that Nrg1 deficiency reduced axonal density near the lesion in both young and mature mice. These observations highlight an essential contribution of Nrg1 to axonal integrity after injury, particularly in the mature brain where regenerative mechanisms are otherwise limited.

### Nrg1 modulates neuroinflammation in an age-dependent manner

Our findings indicate that Nrg1 loss-of-function leads to a neuroinflammatory phenotype in mature mice after CCD, supporting a role for Nrg1 in regulating the brain’s immune response to injury. Specifically, we observed increased IBA1 labeling intensity and a significant expansion of the IBA1-positive area in mature Nrg1-deficient mice, consistent with heightened microglial activation. These changes suggest that, in the absence of Nrg1, microglia adopt a more reactive state, and that Nrg1 normally helps to limit the extent of microglial reactivity following cortical damage.

Importantly, this phenotype was absent in young Nrg1-deficient mice, implying that the anti-inflammatory function of Nrg1 becomes increasingly critical with age. This finding aligns with the view that aging brains rely more heavily on intrinsic neuroprotective mechanisms and suggests that Nrg1 loss may exacerbate maladaptive immune responses in older individuals. Unlike previous studies focusing on exogenous Nrg1 in models of severe injury (Alizadeh et al., 2017; Kataria et al., 2019; Xu et al., 2005), our results highlight the physiological role of endogenous Nrg1 in modulating neuroinflammation even after moderate cortical trauma. Together, these observations extend the functional relevance of Nrg1 beyond axonal regeneration and underscore its contribution to age-sensitive neuroimmune homeostasis.

### Nrg1 controls PNN Integrity and Circuit Plasticity

A particularly novel aspect of our study is the analysis of PNN remodeling after CCD in Nrg1-deficient mice. PNNs are critical regulators of synaptic stability and plasticity, particularly in fast-spiking parvalbumin (PV) interneurons, which express high levels of ErbB4, the main cortical receptor for Nrg1 (del Pino et al., 2013; Fazzari et al., 2010). Our results show a prominent degradation of PNNs in the perilesional cortex after CCD, consistent with previous findings (González-Manteiga et al., 2022). In addition, we observed a further reduction in PNN integrity in the contralesional region, specifically in mature Nrg1-deficient mice. This emphasizes that Nrg1 signaling is essential for maintaining inhibitory network stability and plasticity in the injured cortex.

The degradation of PNNs in mature Nrg1-deficient mice may have several functional implications. First, loss of PNNs around PV interneurons could destabilize inhibitory circuits, leading to an altered excitation/inhibition balance, which is a key determinant of plasticity and recovery (Wang and Fawcett, 2012). Second, impaired PV neuron function could disrupt the synchronization of neuronal assemblies necessary for motor function (Hsieh et al., 2017; Devienne et al., 2021; Wang and Fawcett, 2012). Although the molecular mechanism remains unclear, Nrg1 may regulate PNN maintenance by signaling to Erb4, which is expressed in PV cells. Alternatively, Nrg1 signaling may regulate the PNN through neuron-glia interactions. For instance, Nrg1 may influence the secretion of extracellular matrix remodeling enzymes, such as matrix metalloproteinases (MMPs).

### Implications of Nrg1 for Motor function

The functional deficits observed in motor recovery after CCD in the RPT assay further underscored the importance of Nrg1. The deficits observed in the rotating pole test (RPT) after CCD highlight the functional consequences of Nrg1 loss. Nrg1-deficient mice, particularly mature animals, showed reduced performance compared with controls, consistent with the disruption of callosal connectivity and PNN integrity described above.

These behavioral results support the view that the molecular and cellular alterations caused by Nrg1 deficiency translate into measurable functional deficits. They also reinforce the physiological relevance of Nrg1 signaling in the context of traumatic brain injury, in line with previous studies linking callosal integrity to motor performance.

### Limitations and Future Directions

While our study provides novel insights, a number of limitations must be considered. Mechanistically, how Nrg1 promotes axonal regeneration and supports cortical circuitry remains to be fully elucidated. Nrg1-ICD has been shown to regulate gene expression, although the exact target genes and pathways in the context of injury remain unknown (Bao et al., 2003; Chen et al., 2010). It is plausible that Nrg1-ICD controls gene networks related to axonal growth, synaptic plasticity, or cytoskeletal remodeling, including growth-associated proteins such as GAP43 (see Rodríguez-Prieto et al., 2024). In addition, Nrg1 targets may be involved in inflammation, maintenance of the PNN, and/or neuronal survival. For example, we have previously shown that hypoxia triggers intracellular Nrg1 signaling, which in turn decreases the expression of pro-apoptotic genes (see Navarro-González et al., 2019). Future transcriptomic and proteomic analyses will be instrumental in identifying downstream effectors of Nrg1 signaling during recovery.

Our experiments were performed in the CCD model, which targets the motor cortex. Whether similar mechanisms operate in other cortical or subcortical regions remains to be tested. Moreover, we focused on interhemispheric callosal projections, but other long-range tracts, such as corticospinal or thalamocortical pathways, may also depend on Nrg1 signaling.

### Conclusion

Our study supports a critical role for endogenous Nrg1 and its intracellular signaling in orchestrating the brain’s response to cortical injury. Specifically, we showed that Nrg1 signaling contributes to axonal regeneration, maintenance of cortical wiring, and functional recovery. These effects were particularly evident in the aging brain, suggesting an increased dependence on trophic support with age. Altogether, this work identifies Nrg1 signaling as a promising therapeutic avenue for mitigating the consequences of traumatic brain injury.

## Ethics approval

All animal experiments were conducted in accordance with the guidelines of the Ethical Committee of the Centro de Investigación Príncipe Felipe (CIPF, Valencia, Spain). Animal housing and care were provided by the accredited CIPF Animal Facility. All procedures were approved by the CIPF Bioethics Committee and authorized by the Comunidad Valenciana (Spain), in compliance with European Union Directive 2010/63/EU and Spanish legislation on animal research.

## Acknowledgments

The authors thank the members of the Laboratories of Cortical Circuits in Health and Disease and Neuronal Regeneration at the Centro de Investigación Príncipe Felipe, especially to Drs. Victoria Moreno-Manzano and María Pedraza, for their valuable input, constructive discussions, and continuous support during the course of this study. We are also grateful to Alfredo Collado for his assistance with research materials and logistics, and to the staff of the CIPF Animal Facility for their excellent technical support and animal care. Thanks to Dr. Rosa Viana for her assistance with image acquisition using the Leica Aperio in the IBV microscopy core. The authors used ChatGPT (OpenAI, 2025) as a writing aid to enhance manuscript clarity, consistency, and readability.

## Data availability

All reagents and additional information about the results and methodology are available upon request.

## Abbreviations

AAV: adeno-associated virus
ANOVA: analysis of variance
BSA: bovine serum albumin
CCD: controlled cortical damage
Ctrl: control
DAPI: 4’,6-diamidino-2-phenylindole
ErbB4: erythroblastic oncogene B receptor 4
GFP: green fluorescent protein
IBA1: ionized calcium-binding adaptor molecule 1
KO: knockout
LDA: linear discriminant analysis
NRG1: neuregulin-1
PBS: phosphate-buffered saline
PFA: paraformaldehyde
PNN: perineuronal net
PV: parvalbumin
ROI: region of interest
SEM: standard error of the mean
WFA: Wisteria floribunda agglutinin.

## SUPPLEMENTAL MATERIAL

**Supplementary Fig. S1.**
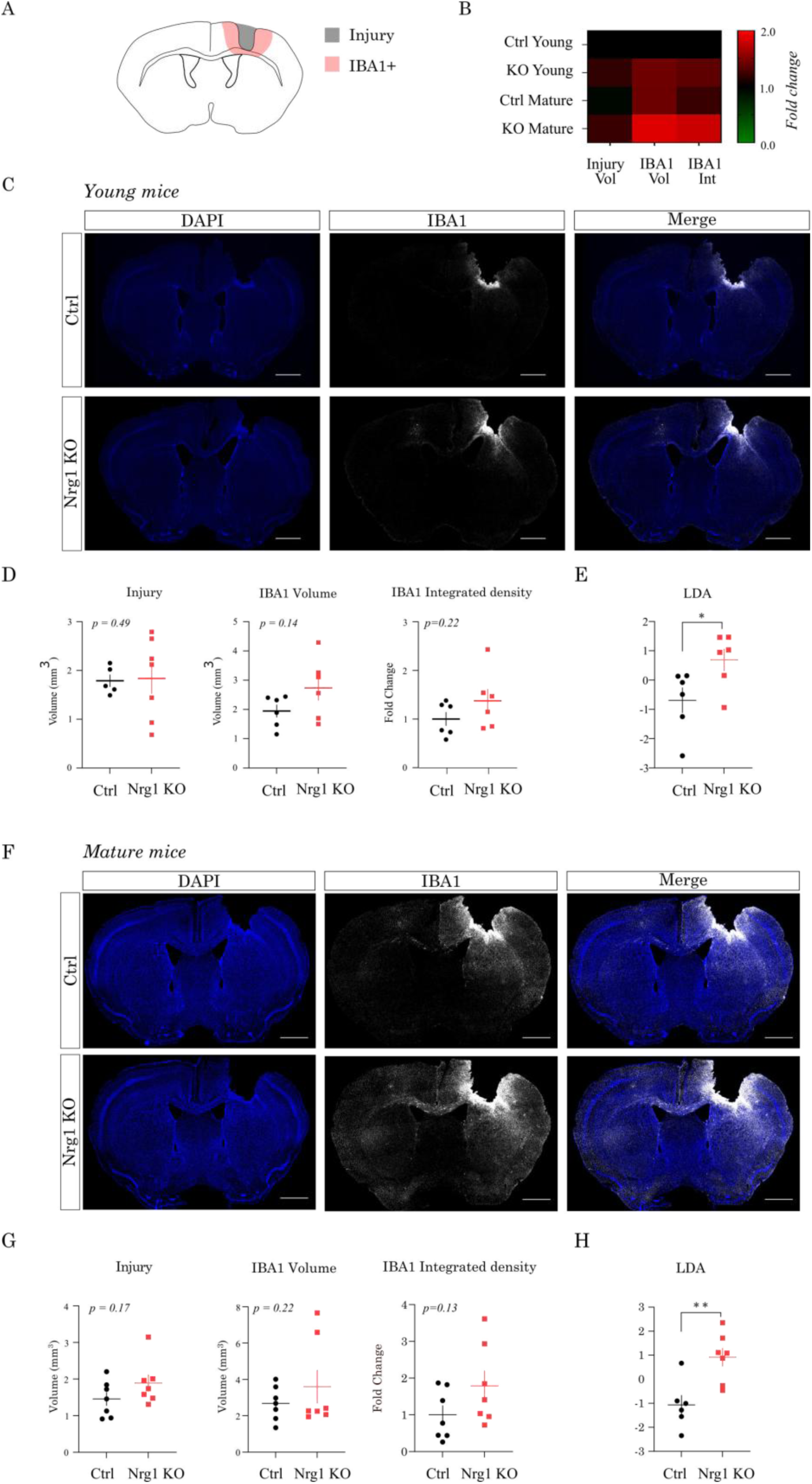
Inflammatory response to CCD in Nrg1-deficient mice. (A) Experimental schema for lesion and inflammation quantification in young (3–4 months) and mature (10–12 months) cohorts, outlining CCD, tissue processing, ROI definition (perilesional cortex and distal reference), and extracted metrics (injury volume, IBA1⁺ volume, IBA1 integrated density). (B) Heatmap of injury and inflammatory metrics per case, grouped by genotype and age; colors indicate relative increases or decreases relative to the control young group. (C) Representative IBA1 immunostaining in perilesional cortex of young mice. IBA1⁺ microglia/macrophages are shown in gray, DAPI in blue. Scale Bar = 500 µm. (D) Univariate quantification of injury volume, IBA1⁺ volume, and IBA1 integrated density in young mice. Graphs show mean ± SEM (Ctrl, n = 6 animals; Nrg1 KO, n = 6 animals). Statistical analysis: unpaired Student’s t-test (p values in graphs). (E) Multivariate linear discriminant analysis (LDA) integrating the same variables as in (D) suggests partial genotype-dependent separation in young mice (Student’s t-test. *p < 0.05). (F) Representative IBA1 immunostaining in perilesional cortex of mature mice. IBA1⁺ microglia/macrophages are shown in gray, DAPI in blue. (G) Univariate quantification of injury volume, IBA1⁺ volume, and IBA1 integrated density in mature mice. Graphs show mean ± SEM (Ctrl, n = 6 animals; Nrg1 KO, n = 6 animals). Statistical analysis: unpaired Student’s t-test (p values in graphs). (H) Multivariate LDA integrating the same variables as in (G) shows clear genotype-dependent separation in mature mice (Student’s t-test. **p < 0.01).

